# Polychrome: Creating and Assessing Qualitative Palettes With Many Colors

**DOI:** 10.1101/303883

**Authors:** Kevin R. Coombes, Guy Brock, Zachary B. Abrams, Lynne V. Abruzzo

## Abstract

Although **R** includes numerous tools for creating color palettes to display continuous data, facilities for displaying categorical data primarily use the **RColorBrewer** package, which is, by default, limited to 12 colors. The **colorspace** package can produce more colors, but it is not immediately clear how to use it to produce colors that can be reliably distingushed in different kinds of plots. However, applications to genomics would be enhanced by the ability to display at least the 24 human chromosomes in distinct colors, as is common in technologies like spectral karyotyping. In this article, we describe the **Polychrome** package, which can be used to construct palettes with at least 24 colors that can be distinguished by most people with normal color vision. **Polychrome** includes a variety of visualization methods allowing users to evaluate the proposed palettes. In addition, we review the history of attempts to construct qualitative color palettes with many colors.

## 1. Introduction

Numerous tools exist in R for creating color palettes that can be used to display continuous data, usually in the form of two-dimensional images or heatmaps. These include the built-in palettes (`rainbow, heat.colors, terrain.colors, topo.colors, and cm.colors`) as well as the colorRamp function in the **grDevices** package. Using `colorRamp`, researchers can construct arbitrary sequential palettes (to emphasize the change from low to high values) or divergent palettes (to emphasize the difference between large negative values, values near zero, and large positive values). The options for creating qualitative palettes to display discrete categorical data in R, however, are more limited. The primary source of qualitative palettes is the **RColorBrewer** package (Neuwirth 2014). In addition to sequential and divergent palettes, **RColorBrewer** contains eight discrete palettes. These palettes were designed primarily with a view toward coloring map regions (Brewer 1989; Harrower and Brewer 2003), and thus a major design consideration was how easily colors could be distinguished when two large blocks of color appeared next to each other. These palettes may not work as well for other common kinds of plots: bar charts, scatter plots, line plots, or others where recognizability against a different background becomes more important. Two of the Brewer palettes consist of pastels, which are most suitable for display against a dark background. Four contain darker colors, more suitable for display against a white background. The remaining two contain a mix of both lighter and darker shades. As a result, for most kinds of plots, one must choose from a very small number of qualitative palettes. For a more complete discussion of the properties and roles of sequential, diverging, and qualitative palettes, see Zeileis, Hornik, and Murrell (2009).

In addition, a critical limitation is that the **RColorBrewer** palettes contain at most 12 distinct colors. While adequate for many applications, this limitation presents potential roadblocks whenever a researcher tries to present data that highlight a categorical variable or variables with more than 12 distinct values. The particular application that caused us to confront this limitation arose from the increasingly common biologically derived datasets produced by “omics” scale technologies. In this context, it is natural to use color in plots to indicate the chromosome of origin of the data. Because there are 22 human autosomal chromosomes plus two sex chromosomes, this application requires 24 distinct colors.

Interestingly, some palettes are already in use for chromosome data. The cytogenetic technique of spectral karyotyping (SKY; Garini, Macville, du Manoir, Buckwald, Lavi, Katzir, Wine, Bar-Am, Schröck, Cabib, and Ried (1996)) uses mixtures of fluorescent dyes to label each chromosome differently. Because several of the dyes actually fluoresce in the near-infrared spectrum, the resulting images are conventionally presented on a black background, using false color mappings to display the chromosomes with 24 different colors. However, the most common false color palettes almost always include some pairs of colors that are difficult to distinguish reliably. This weakness is less important for SKY, since cytogeneticists can often use the size and other characteristics of chromosomes to tell them apart. But the same palettes are unlikely to work as well in a scatter plot.

In this manuscript, we describe the **Polychrome** package, which contains methods to create “large” palettes with up to 40 distinct colors that can be reliably distinguished by most (noncolor-blind) individuals. The algorithm is inspired by (relatively old) experimental data on how well-separated colors must be in order for a typical viewer to tell them apart. The package also contains several historical large color palettes, a variety of tools to illustrate how palettes might work with different kinds of plots, and a set of tools to assign consistent color names to the resulting members of each palette. The package is available from CRAN. Source code is available through R-Forge as one of the Object-Oriented Microarray and Proteomics Analysis (OOMPA) packages described at http://oompa.r-forge.r-project.org/. The latest version be can also installed from the official OOMPA repository as follows:

~~~
*R> source(“http://silicovore.com/OOMPA/oompaLite.R“)
R> oompainstall(“Polychrome”)*
~~~

### 1.1. Color spaces

Throughout this manuscript, we will use several color spaces. A color space is an organized system of colors that, when combined with a description of a physical device, allows for reproducible representations of color. Table 1 lists some common color spaces. (An accessible description of color spaces and mathematical transforms between them can be found at https://en.wikipedia.org/wiki/Color_model.) The most common color space for computer monitors is the standard red-green-blue (sRGB) color space. This color space relies on the additive color properties of light and, to be fully specified, requires a definition of a reference “white point” and a “gamma” correction (*γ* = 2.2 in the standard space) in addition to the three primary colors. When mixing pigments for printing, one needs a subtractive color model such as CMYK. Here the issue is that the observed color when illuminating an object depends on which wavelengths are absorbed (or subtracted) and which wavelengths are reflected. Among the pigment primary colors, cyan (C) absorbs red and reflects blue and green; magenta (M) absorbs green and reflects red and blue; yellow (Y) absorbs blue and reflects red and green. Black (K) is added to increase the overall absorbance and thus decrease the overall luminance.

**Table 1:** Common color spaces. (R = red, G = green, B = blue; C = cyan, M = magenta, Y = yellow, K = black; H = hue, S = saturation, V = value, L = luminance or lightness; CIE = International Commission on Illumination).

| Color space | Context | Notes |
| --- | --- | --- |
| sRGB | light | standard red-green-blue |
| CMYK | pigment | cyan, magenta, yellow, black |
| HSV | cylindrical | hue, saturation, value |
| HSL | cylindrical | hue, saturation, luminance |
| CIE $L^*u^*v^*$ | light | |
| CIE $L^*a^*b^*$ | pigment | |
| HCL | cylindrical | hue, chroma, luminance |

Many artists prefer to think in the “cylindrical” color spaces HSV or HSL. In these spaces, the hue (H) is given by the angle around a central axis, starting with red at 0°, green at 120°, and blue at 240°. Saturation (S) is the colorfulness of a stimulus relative to its background; it is represented by the distance from the central axis. In the HSL color space, distance along the central axis represents the luminance (L) in candela per square meter relative to the luminance of the reference white point. In the HSV model, the central axis represents the value (V) or lightness, which is the brightness relative to a similarly illuminated white. Because these color spaces have a lack of perceptual uniformity, they will be of limited utility to our project.

The International Commission on Ilumination (CIE) has defined two color spaces in which distances try to more closely reflect perceptual uniformity. The L*u*v* color space is more suited for light-based additive colors, while the L*a*b* color space is more suited for pigment-based subtractive colors. In both spaces, L represents the lightness and the other two axes represent contrasts of “opposing colors”. The a* axis goes from green to magenta, and the b* axis goes from blue to yellow. The u* axis goes from purple to yellow, and the v* axis from cyan to red. The HCL color space is another cylindrical model that uses chroma instead of saturation; chroma is an absolute measure of colorfulness, where saturation is relative to the maximum colorfulness for a given hue and value. In most cases, HCL can be viewed as a cylindrical transformation of L*u*v* space.

## 2. A brief history of large color palettes

One of the earliest discussions of a large qualitative palette was the system of 22 colors described by Kenneth Kelly (Kelly 1965). At the time, Kelly worked for the United States National Bureau of Standards (now called the National Institute of Standards and Technology) and was responding to multiple requests for high-contrast palettes of various sizes. The first nine colors (starting with “white” and proceeding through “gray”) in his proposed palette agree with an earlier list produced by Judd (Judd 1949) and should provide maximum contrast even for viewers who are red-green color blind. Each of the other colors (10–22) “has been selected so that it will contrast maximally with the color just preceeding it and satisfactorily with the earlier colors”. Kelly did not describe any numerical criteria to explain the terms “maximally” nor “satisfactorily”.

Kelly’s set of 22 colors is made available in the **Polychrome** package through the function kelly.colors. The assigned color names are taken from his paper.

~~~
*R> library(“Polychrome”)
R> Kelly <-kelly.colors(22)*
~~~

Using the `swatch` function from **Polychrome**, the resulting palette is displayed in Figure 1.

**Figure 1:**
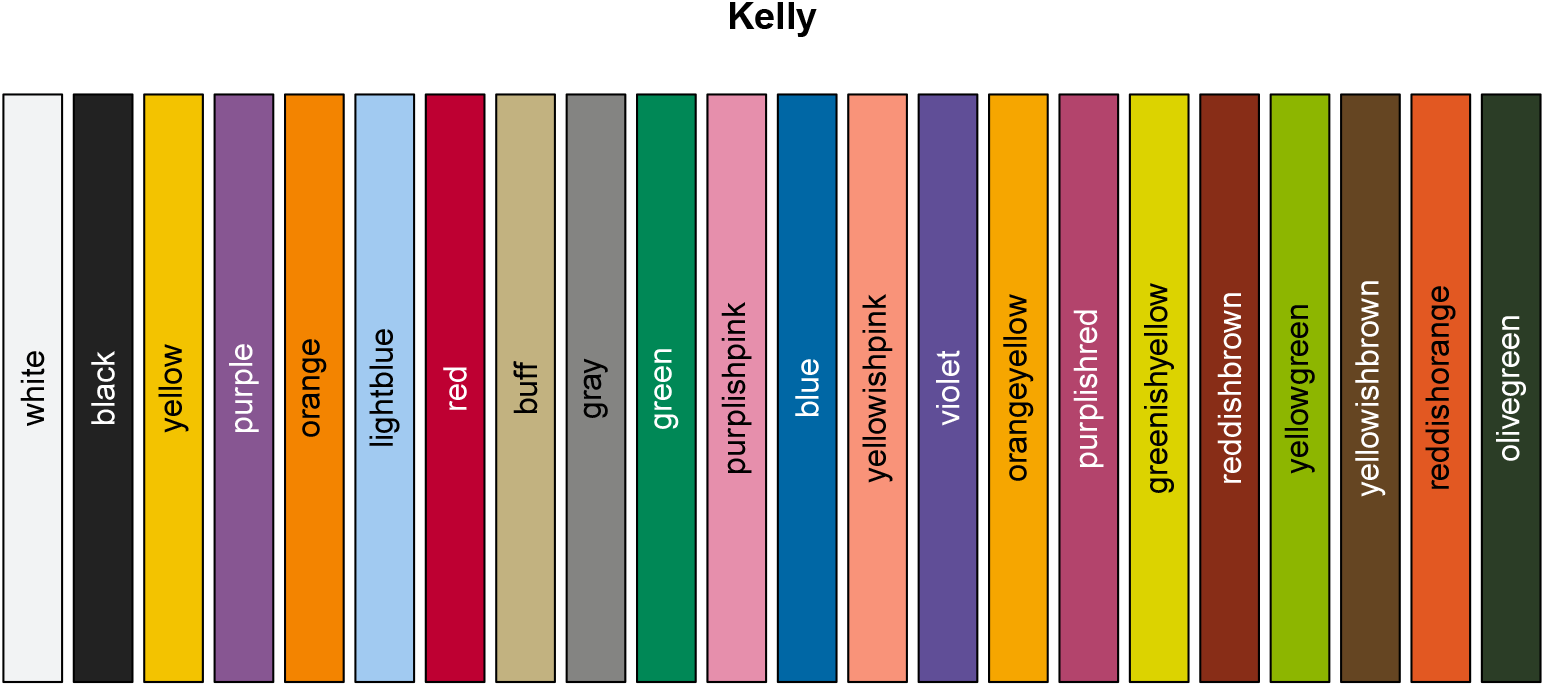
The 22-color qualitative palette defined by Kelly in 1965.

~~~
*R> swatch(Kelly)*
~~~

### 2.1. Quantitative construction of qualitative palettes

The first quantitative approach to constructing large qualitative palettes was presented by Carter & Carter (Carter and Carter 1981, 1982). They made two critical observations:

1. The ability of a typical color-normal human observer to distinquish two colors appears to be monotonically related to the Euclidean distance between their coordinates in L*u*v* space, as defined by the CIE.
2. Based on experimental observations, the minimum distance at which two colors can be reliably distinguished is about 40 CIE L*u*v* units.

#### Distinguishability of the Kelly colors

Interestingly, the Kelly palette does not satisfy the criterion derived by Carter & Carter (Figure 2). In fact, only 13 colors can be selected from his set of 22 colors to keep the minimun distance between them at least 40. The plot in Figure 2 was made using the `plotDistances` function from **Polychrome**. This function reorders colors in a palette to put the most distinguishable colors first. It then plots the minimum distance of the *n*^th^ color from the previous *n* — 1 colors.

**Figure 2:**
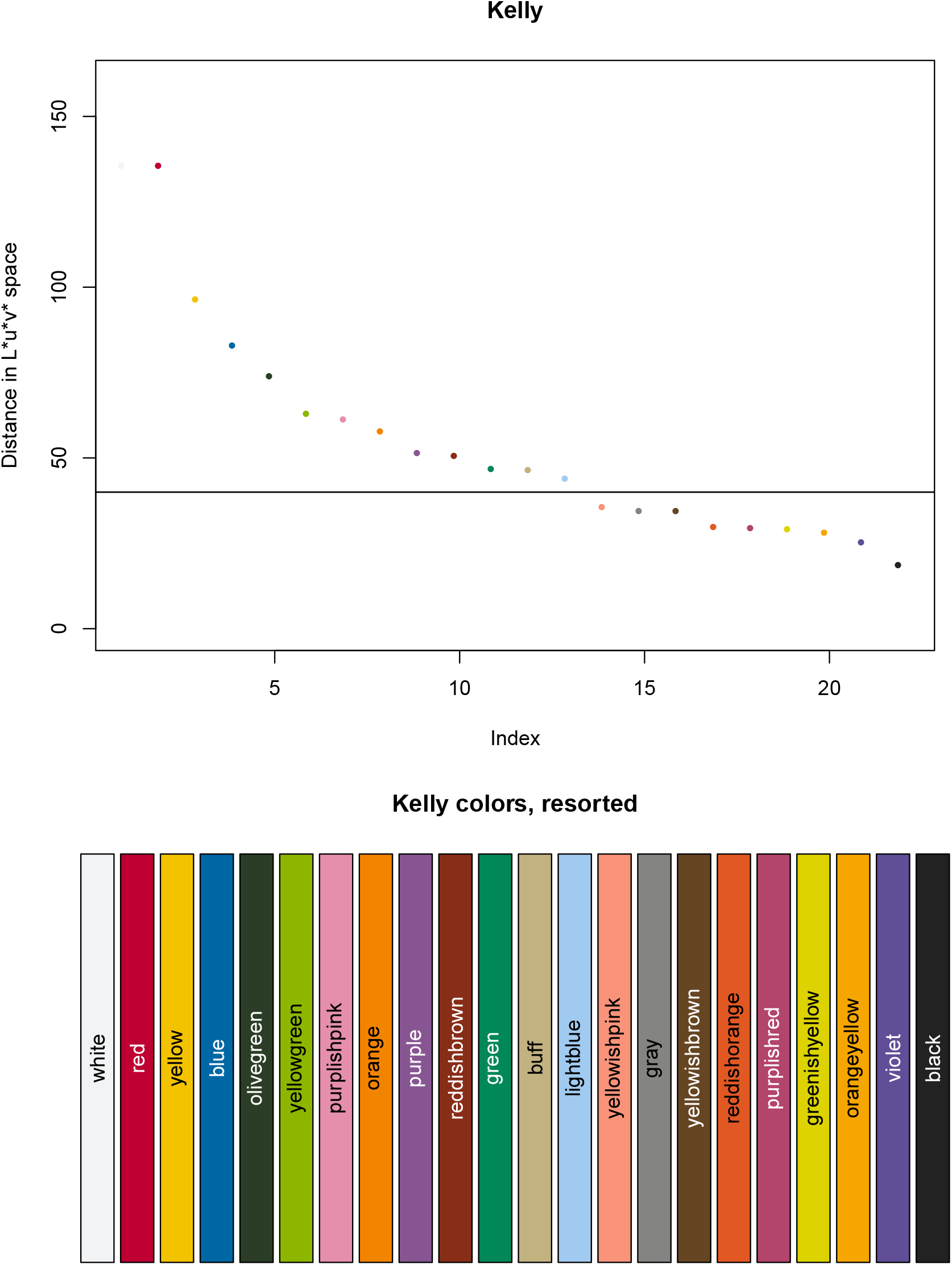
(Top) Minimum relative distance between Kelly colors. (Bottom) Kelly colors sorted to maximize the contrast at each step between the next color and all preceeding colors.

~~~
*R> layout(1:2, widths = 1, heights = c(3, 2))
R> opar <− par(mai = c(0.82, 0.82, 0.82, 0.42))
R> ks <− plotDistances(Kelly, pch = 16, cex = 2, ylim=c(0, 160))
R> abline(h = 40)
R> par(opar)
R> swatch(ks$colors, main = “Kelly colors, resorted”)
R> layout(1, 1, 1)*
~~~

#### Qualitative rainbow palettes

Ihaka suggested that qualitative palettes could be constructed in hue-chroma-luminance (HCL) space by holding the chroma and luminance constant and spreading the hue out uniformly across the spectrum (Ihaka 2003). This idea has been implemented in the `rainbow_hcl` function in the **colorspace** package (Ihaka, Murrell, Hornik, Fisher, and Zeileis 2016). These rainbow color schemes work very well with small palettes containing four to six colors, as illustrated in Ihaka’s paper. Here, however, we construct a 24-color rainbow palette.

~~~
*R> library(“colorspace”)
R> Rainbow <− rainbow_hcl(24, c = 90, l = 50)*
~~~

We plotted the rainbow palette in Figure 3. Unfortunately, only nine of the rainbow colors can be reliably distinguished by most observers. (The `computeDistances` function determines the minimum distance of each color to all preceding colors in the palette; we can use this to detemine how many distances are large enough.) As with the palettes produced by **RColorBrewer**, we would still expect these palettes to be useful when coloring maps or in other applications that rely on broad areas of color.

**Figure 3:**
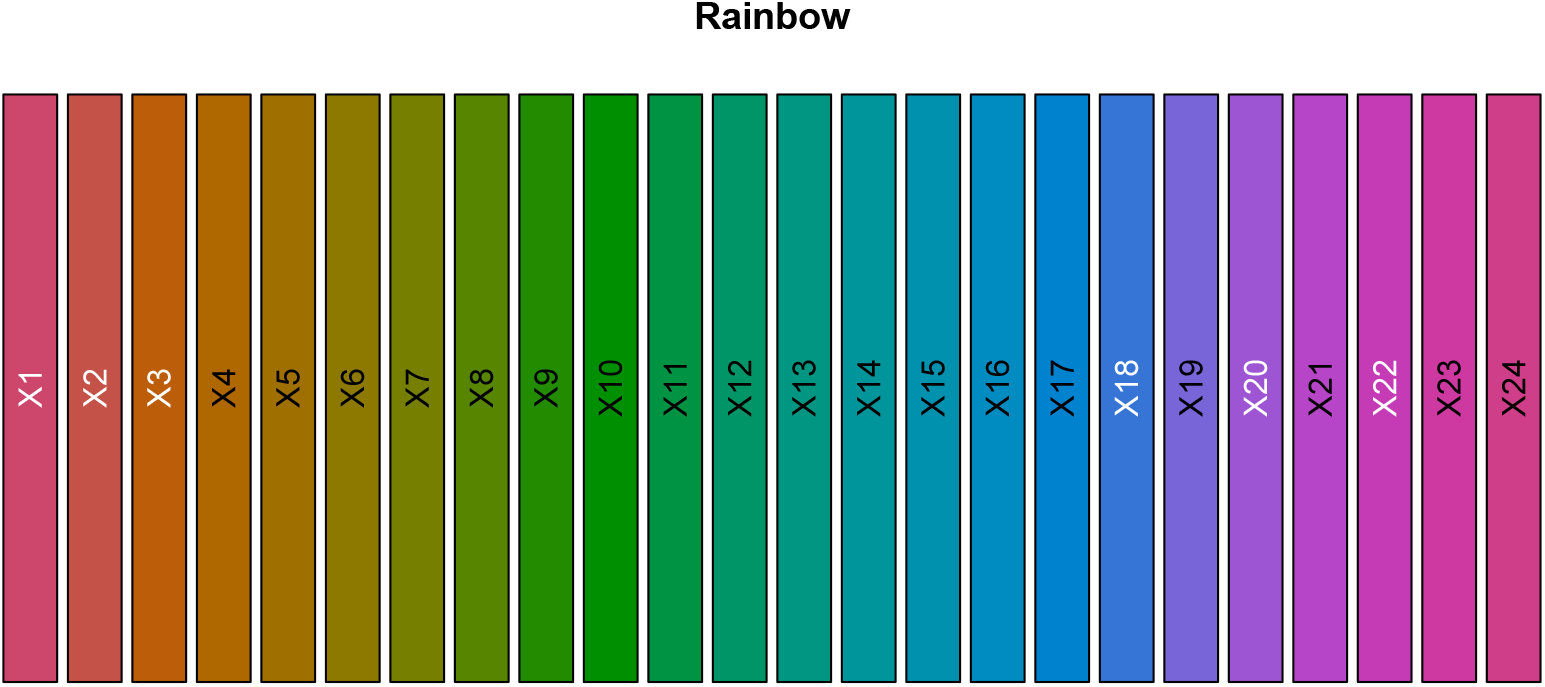
A 24-color rainbow palette.

~~~
*R> rd <− computeDistances(Rainbow)
R> sum(rd > 39)*
~~~

~~~
[1] 9
~~~

~~~
*R> swatch(Rainbow)*
~~~

#### The Carter and Carter algorithm

Given the two examples we have looked at so far, one is led to wonder if it is even possible to construct a qualitative palette with 24 distinguishable colors. Carter & Carter point out that the problem of finding an optimal set of *N* colors is equivalent to placing *N* points into the (displayable) region of CIE L*u*v* space in such a way as to maximize the minimum distance *D* between any pair of points. They describe a heuristic algorithm for solving the problem. Briefly, you start with an initial step size *S* and randomly select *N* points. At each iteration, you find the pair of points separated by the current minimal distance *D*. Evaluate all possible moves consisting of changing one coordinate by *S* units (either positively or negatively). Make the move that most increases *D*. Then halve the step size and repeat. To avoid converging to local optima, they add an extra step to start over again at the initial step size after you reach the resolution of the target device. The algorithm halts when no moves that increase *D* are possible. They reported that running the algorithm multiple times with *N* = 25 colors yielded palettes for which the mean *E*[*D*] = 51.60 and the variance *V*[*D*] = 7.34. Based on these results, they also speculated that *N* = 25 might be near the maximum size of a reliably distinguishable qualitative color palette. For our purposes, unfortunately, they did not provide a specific example of a 25-color palette; the only example they gave was an optimal six-color palette.

### 2.2. A 32-color palette

Working in CIE L*a*b* color space instead of CIE L*u*v* space, Glasbey and colleagues constructed a well-separated 32-color palette (Glasbey, van der Heijden, Toh, and Gray 2007). They used a greedy sequential search algorithm. Given an initial set of colors, they searched through all 256^3^ colors in an RGB cube to find the color that maximized the minimum distance (in L*a*b* space) to the current set of colors. Adding this color to the set, they iterated the procedure until they obtained the desired number of colors. Recognizing that this algorithm did not guarantee an optimal solution, they also described a modification using simulated annealing. Bianco and Citrolo described a similar approach using greedy algorithms or simulated annealing, and added the idea of using a genetic algorithm (Bianco and Citrolo 2013); however, they did not provide an explicit example of a large palette.

We have included the Glasbey palette in the **Polychrome** package; it is accessible via the function `glasbey.colors`.

~~~
*R> Glasbey <− glasbey.colors(32)*
~~~

This palette is displayed in Figure 4. Depending on how strictly one interprets the Carter & Carter criterion, about 28 of the Glasbey colors should be reliably distinguishable by a color-normal observer.

**Figure 4:**
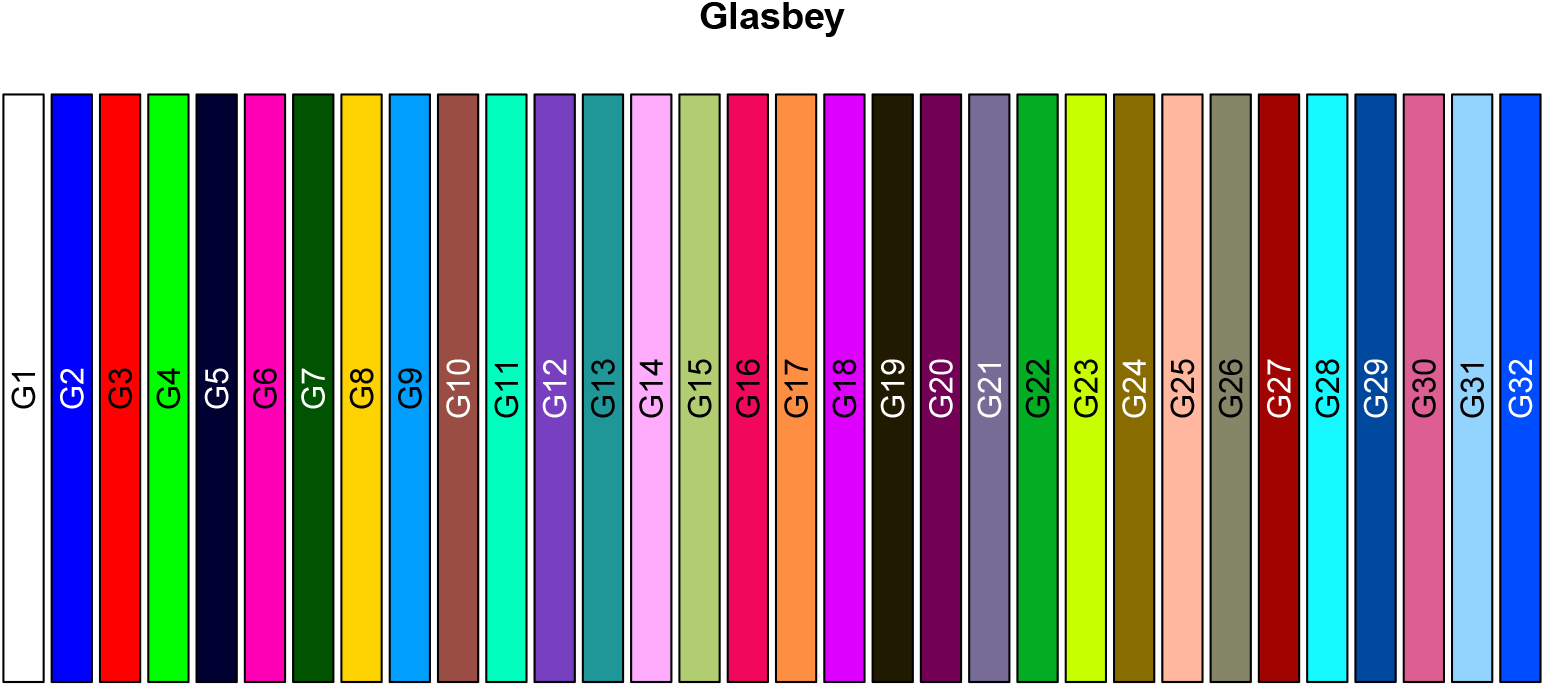
The 32-color Glasbey palette.

~~~
*R> gd <− computeDistances(Glasbey)
R> sum(gd > 39)*
~~~

~~~
[1] 28
~~~

~~~
*R> swatch(Glasbey)*
~~~

### 2.3. An alphabet of colors

Large qualitative color palettes were revisited by Green-Armytage (2010). The impetus for this study was a claim by Rudolph Arnheim that an alphabet composed of 26 colors instead of shapes would be unusable (Arnheim 1974). A workshop for the members of the Colour Society of Australia in 2007 took up the challenge of constructing a “color alphabet” and compared it to alphabets made of symbols. They found that this color alphabet could be read more easily than one made up of abstract symbols from the “Dingbats” font. A follow-up study compared an alphabet made of “nameable shapes” to one made of “nameable colors”.

The names in both cases were chosen so that there was one shape or color beginning with each letter of the alphabet. In this test, deciphering text using the shapes was slightly faster on average (2 min 55 sec) than using colors (3 min 20 sec). Nevertheless, the experiments did provide solid evidence that it was not only possible to construct a palette with 26 different reproducibly distinguishable colors, but to use that palette as an alphabet.

The final Green-Armytage “color alphabet” is included in the **Polychrome** package. Users can access it with the function `green.armytage.colors`. The color alphabet is displayed in Figure 5. Interestingly, this palette was found to be usable even though only a subset of 17 colors meets the criterion of Carter & Carter, and four pairs of colors are separated by distances less than 25.

**Figure 5:**
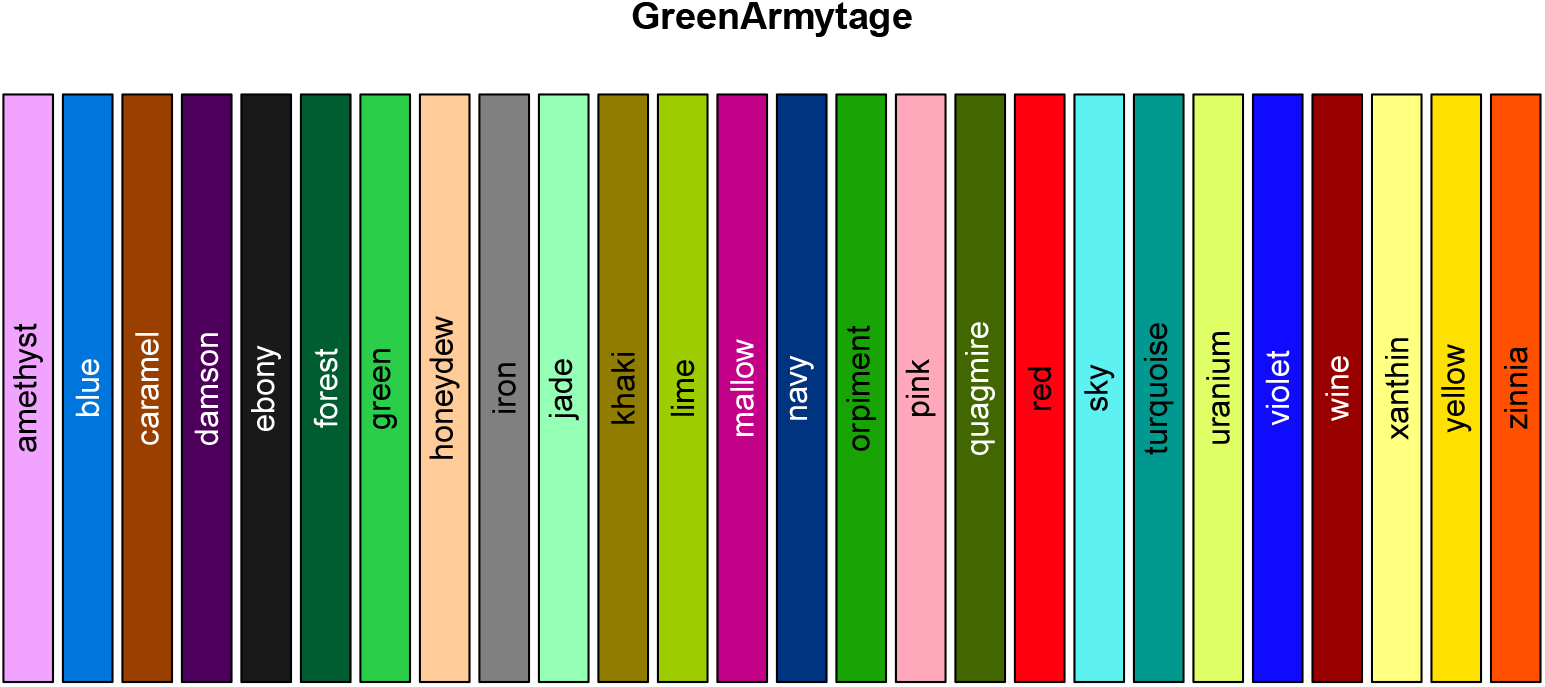
The 26-color Green-Armytage alphabet palette.

~~~
*R> GreenArmytage <− green.armytage.colors(26)
R> gad <− computeDistances(GreenArmytage)
R> sum(gad > 39)*
~~~

~~~
[1] 17
~~~

~~~
*R> sum(gad < 25)*
~~~

~~~
[1] 4
~~~

~~~
*R> swatch(GreenArmytage)*
~~~

## 3. The Polychrome algorithm to create palettes

Our goal, frankly, is not to find optimally separated sets of *N* colors for arbitrary N. We simply want to find sets with *N* ≈ 30 that are “good enough”, where the definition of “good enough” is that it meets the Carter & Carter criterion that the minimum distance in CIE L*u*v* space is at least 40. Toward that end, we have implemented a variant of the greedy algorithm described in Glasbey *et al.* (2007). Specifically, we start by randomly selecting a set *S* containing a large number (by default, M = 50,000) of points in CIE L*u*v* space. The user seeds the algorithm with one or more colors that they would like included in the palette. We then iteratively select the color in *S* that maximizes the minimum distance to the current set of colors; add it, and repeat. The implementation of the algorithm makes heavy use of facilities provided by the **colorspace** package for translating between different color models. The interface to all **Polychrome** functions, however, assumes that colors are written in the compact hexadecimal notation (e.g., “`#A34CD2`”).

## 3.1. A 36-color palette

We used **Polychrome** to create a palette with 36 colors. Since the algorithm requires that we seed it with starting colors, we followed some basic principles of design: “The first color is white. The second color is black. The third color is red” (Black 1997). Specifically, we used the following code, where the three seed colors are adjusted to lie in the desired luminance range:

~~~
*R> set.seed(567629)
R> P36 <− createPalette(36, c(“#474747”, “#E2E2E2”, “#CC0000”))*
~~~

The random seed was set to ensure that the computation was reproducible. We then saved the palette as part of the package; users can now access it using the `palette36.colors` function.

~~~
*R> P36 <− palette36.colors(36)*
~~~

We display the palette along with the plot of minimum distance in Figure 6. All 36 colors should be distinguishable since the minimum distance between pairs of colors is close to 40.

**Figure 6:**
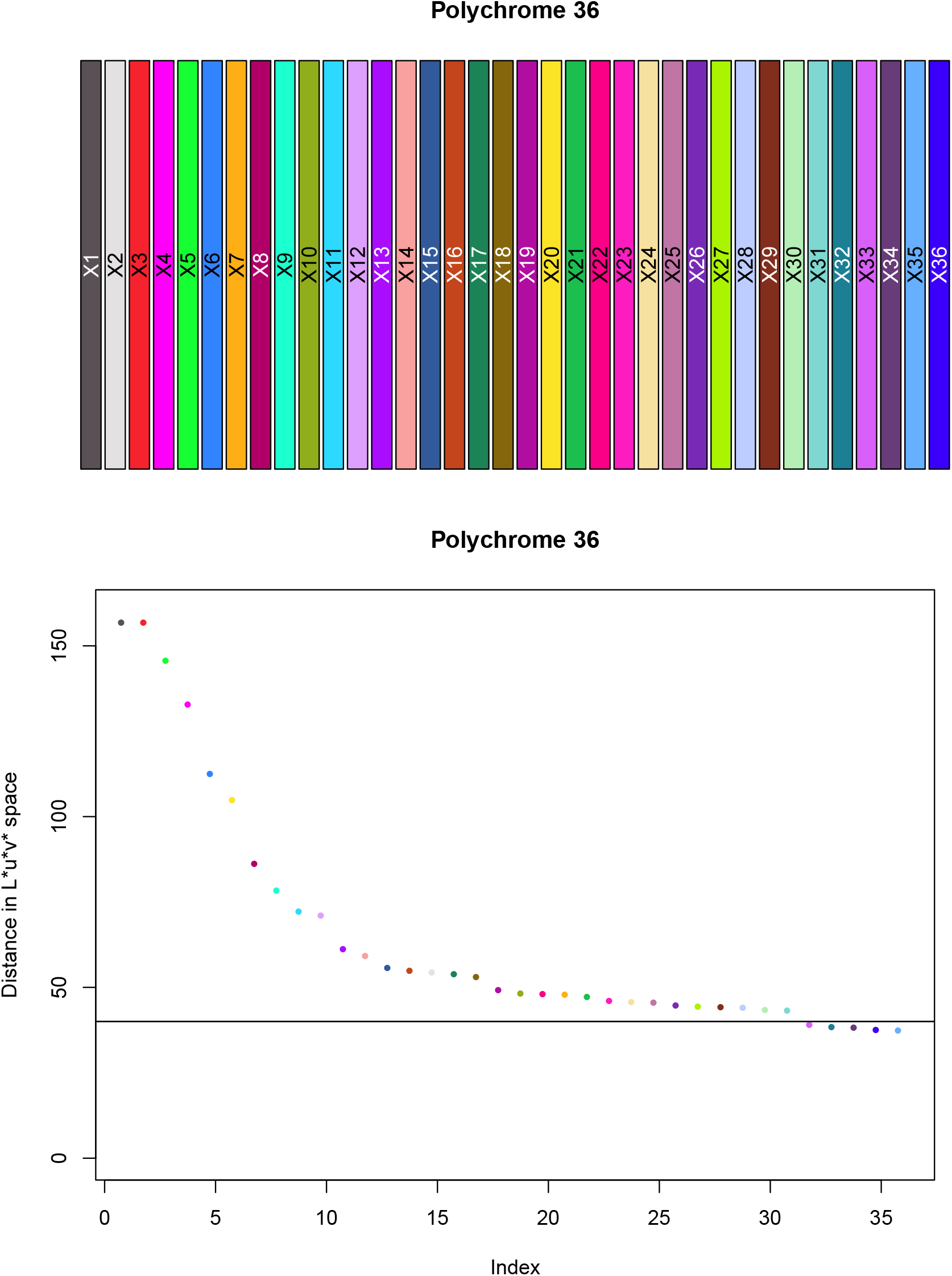
(Top) The 36-color Polychrome palette. (Bottom) The minimum relative distance between colors.

~~~
*R> layout(1:2, widths = c(1, 1), heights = c(2, 3))
R> swatch(P36, main = “Polychrome 36”)
R> opar <− par(mai = c(0.82, 0.82, 0.82, 0.42))
R> pd <− plotDistances(P36, pch = 16, cex = 2, ylim = c(0, 160),
+ main = “Polychrome 36”)
R> abline(h = 40)
R> par(opar)
R> layout(1, 1, 1)*
~~~

### 3.2. A revised alphabet palette

Our 36-color palette is larger than the number of letters in the English alphabet. Moreover, the first 26 colors should be substantially easier to distinguish (since they are well separated in CIE L*u*v* space) than the color alphabet devised by Green-Armytage. So, we created our own “alphabet palette” and included it in **Polychrome** (Figure 7). Following Green-Armytage’s lead, we manually assigned names to each color that start with different letters of the alphabet.

**Figure 7:**
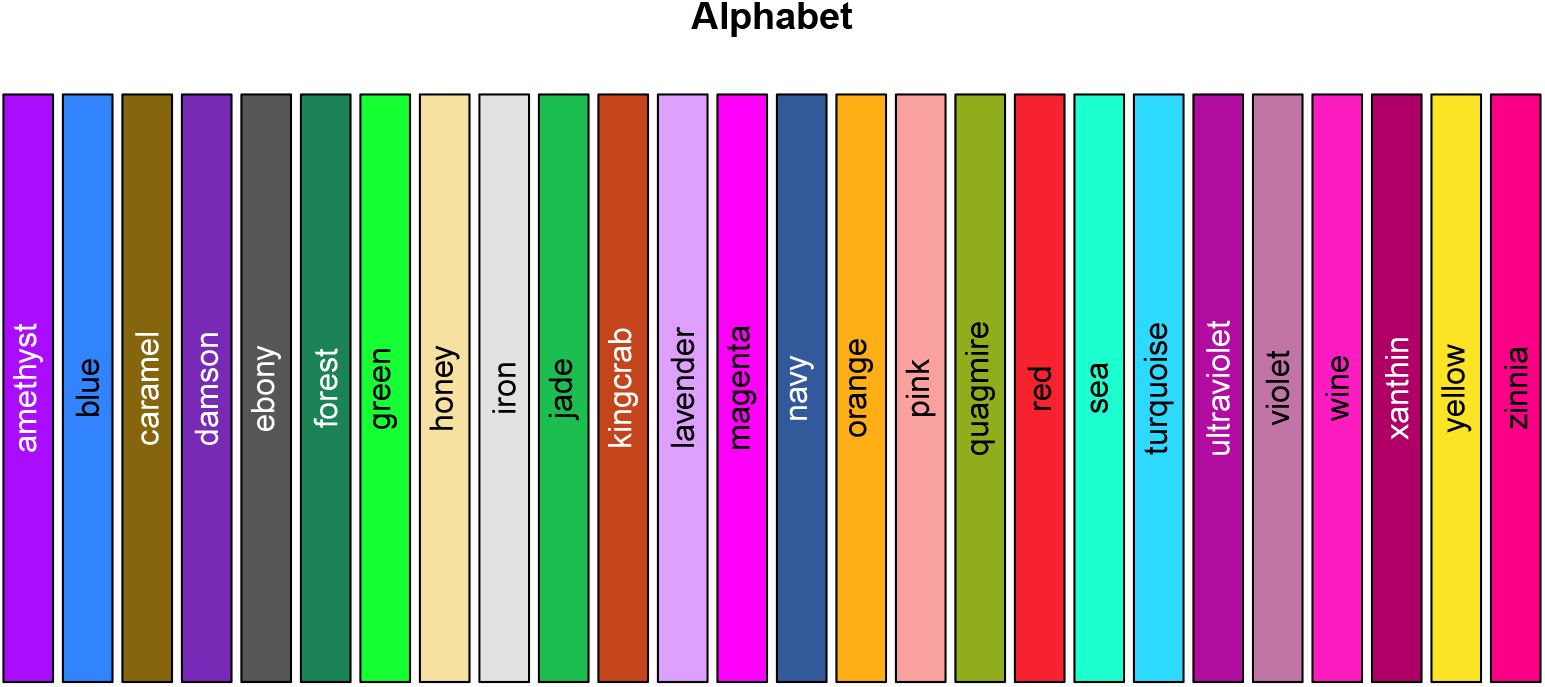
The 26-color qualitative “alphabet” palette derived from **Polychrome** 36.

~~~
*R> Alphabet <− alphabet.colors(26)
R> swatch(Alphabet)*
~~~

## 4. Polychrome methods to display palettes

The only tool we have described so far to assess the usability of qualitative palettes is the `plotDistances` function. However, we believe that it is useful to be able to see the palettes in action in order to decide if they will work for any particular application. Toward that end, we have included a set of visualization functions in the package. In Figure 8, we illustrate the “Polychrome 36” palette using four of these visualization tools:

1. The `uvscatter` function plots one point for each color in the palette using its u-v chromatic coordinates.
2. The `luminance` function plots one point for each color to display its luminance value, in increasing order.
3. The `rancurves` function generates one sine curve, with random phase and amplitude, in each color.
4. The `ranpoints` function generates and plots *N* points in each color, where points of the same color are roughly clustered together.

**Figure 8:**
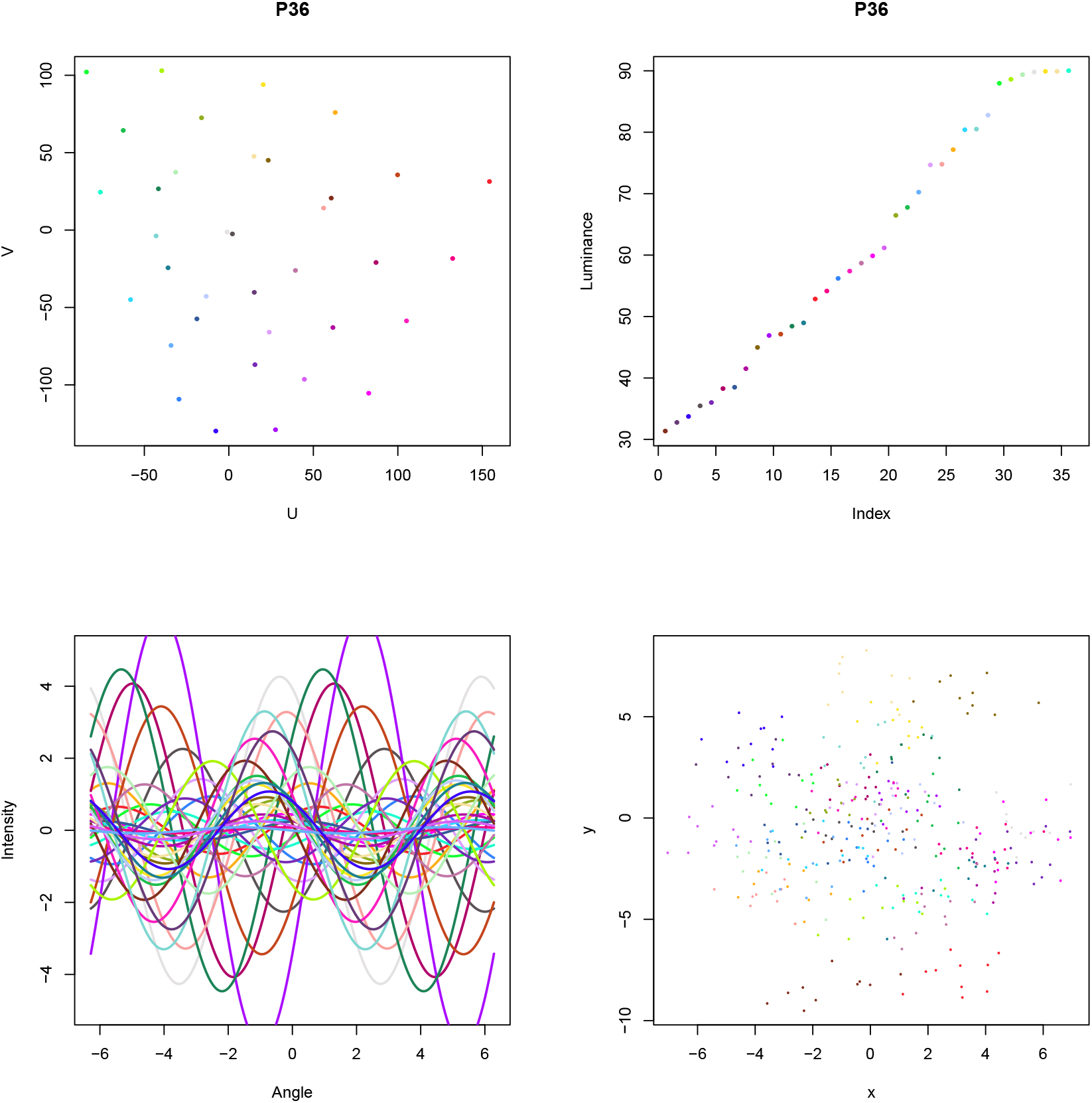
Illustration of the behavior of the **Polychrome** 36 palette. (Top left) Colors displayed by their U-V chromatic coordinates. (Top Right) Sorted display of colors by luminance. (Bottom Left) Random sine curves, one per color. (Bottom Right) Random points, ten per color.

Additional functions (not illustrated here) allow the user to produce bar plots of colors, sorted either by hue or by luminance or in random order. These functions allow the user to confirm that adjacent blocks of color can be distinguished. There are also functions that use the Euclidean distance in CIE L*u*v* space to perform hierarchical clustering or principal components analysis on the colors and then plot the result. These functions give the user alternative views of whether the colors are well separated.

~~~
*R> set.seed(342587)
R> opar <− par(mfrow = c(2, 2), mai = c(0.82, 0.82, 0.82, 0.42))
R> uvscatter(P36)
R> luminance(P36)
R> rancurves(P36)
R> ranpoints(P36)
R> par(opar)*
~~~

### 4.1. Bounding the luminance for various background colors

Both the Kelly 22-color palette and the Glasbey 32-color palette include white and black in their sets of colors. In a statistical application where we want to create scatter plots, one of those colors would almost certainly be used as the background color. As a result, the effective number of colors usable to distinguish levels of a categorical factor must be reduced by at least one. It is possible that very light or very dark colors might also be ruled out depending on the choice of background color. This factor is likely to be even more important when plotting thin lines or small points, when colors will be harder to identify.

To address this issue, the `createPalette` function accepts an optional argument to bound the range of the luminance values. The default value of the `range` parameter, c(30, 90) avoids any colors that are too dark (*L* < 30) or too light (*L* > 90). The “Polychrome 36” palette used the default values, and so should be usable against either a white or black background. However, one can adjust these values to create palettes that are likely to offer better contrast when the background is known.

#### A dark palette for light backgrounds

We used the following code to create a palette where all of the colors are relatively dark. We have confirmed that the minimum distance between colors in the palette is near 40. The colors are displayed in Figure 9.

**Figure 9:**
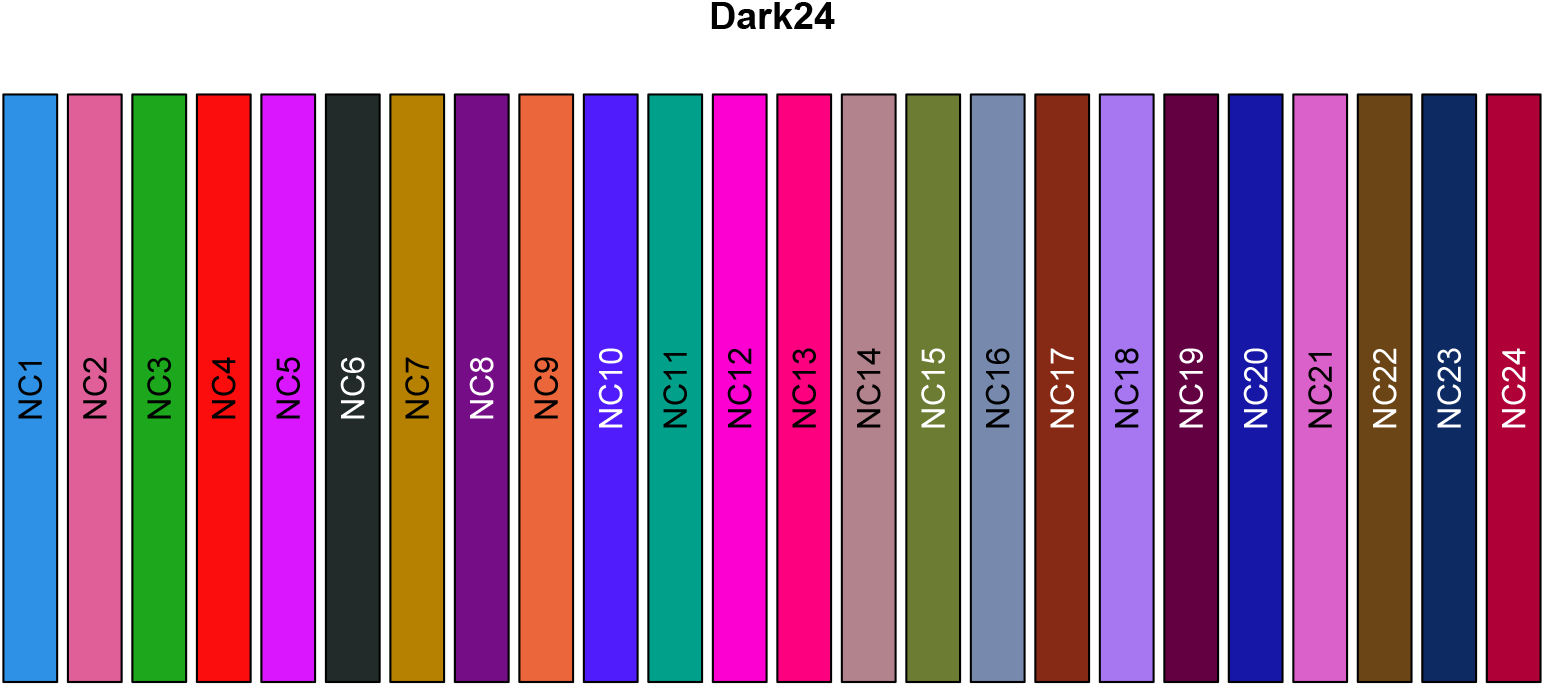
A 24-color palette of well-separated darker colors.

~~~
*R> set.seed(9641)
R> Dark24 <− createPalette(24, c(“#2A95E8”, “#E5629C”), range = c(10, 60),
+ M = 100000)
R> dd <− computeDistances(Dark24)
R> min(dd)*
~~~

~~~
[1] 38.91386
~~~

~~~
*R> swatch(Dark24)*
~~~

#### A light palette for dark backgrounds

We used the following code to create a palette where all of the colors are relatively light. We have confirmed that the minimum distance between colors in the palette is ≥ 40. The colors are displayed in Figure 10. Because we expect this palette to be used against a dark background, the plot has `inverted` the usual foreground and background colors using the invertColors function from `Polychrome`.

**Figure 10:**
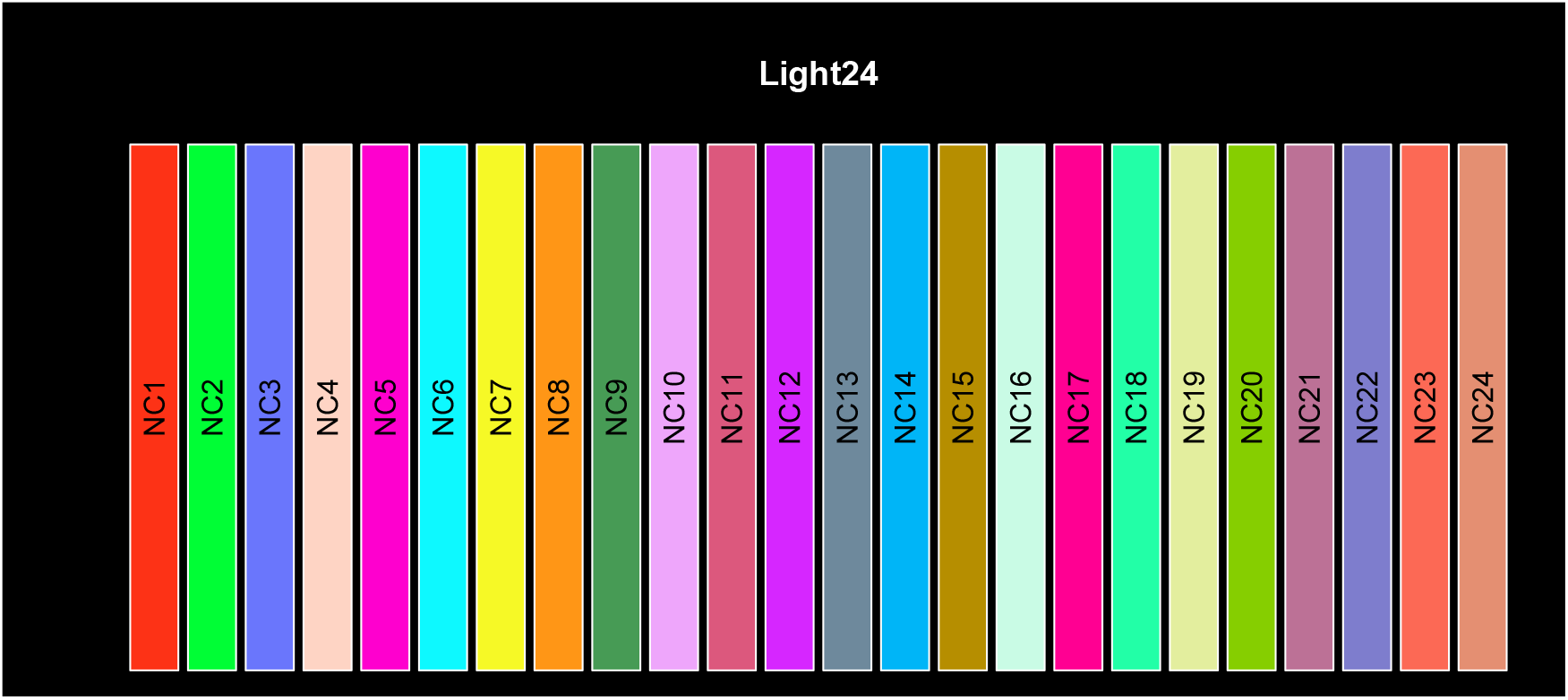
A 24-color palette of well-separated lighter colors.

~~~
*R> set.seed(701138)
R> Light24 <− createPalette(24, “#ff0000”, range = c(55, 95), M = 100000)
R> ld <− computeDistances(Light24)
R> min(ld)*
~~~

~~~
[1] 43.03332
~~~

~~~
*R> opar <− invertColors()
R> swatch(Light24)
R> par(opar)*
~~~

## 5. Color names

One of the lessons to be learned from the Green-Armytage color alphabet is that colors can be used more effectively if one can associate meaningful names to them. Of course, this observation did not originate with Green-Armytage (2010). One can see evidence of it in the classic book by Berlin and Kay (1969), who argue that the underlying biological properties of vision led languages to add color terms in a specific order. They agree with the designers that colors (or at least their names) start with white, black, and red—folowed by green or yellow. A cross-cultural study by Boynton (1989) found eleven colors that are never confused by people with normal color vision: black, white, red, green, yellow, blue, brown, purple, pink, orange, and gray.

Once we get beyond those eleven basic colors, naming becomes rather more complex. The situation is probably not helped by marketers and fashion designers who invent new color names for the “color of the year”. Few people can probably identify 1985’s color called “Kennett Square Pennsylvania”, although it might help if they knew that the city bills itself as the mushroom capital of the world.

The **Polychrome** package draws on two longer and more standard sources of color names that can be assigned to palette members: the ISCC-NBS standard and the UNIX/X11 color list.

The first list of names was produced by a joint effort of the Inter-Society Color Council (ISCC) and the National Bureau of Standards (NBS). They divide color space into a nested or layered collection of contiguous regions and assign names to the centroid of each region. The layers are consistent with the more elaborate system developed by Munsell (1992). There are 13 colors at the first layer, 29 at the second, 267 at the third, and thousands at the fourth. The ISCC-NBS set of color names applies to the 267 colors at the third layer. Using the hexadecimal representations available at http://tx4.us/nbs-iscc.htm, we have built a function, `isccNames`, that assigns the name of the nearest centroid (in CIE L*u*v*) to any hexadecimal color specificaiton.

The second list of names is already available in `R` through the `colors` function in **grDevices**. In Version 3.4.0, this function produces a list of 657 (redundant) color names. The origin of the list goes back to the “colors” file included on early UNIX computers running the X11 windowing system. We have implemented another function, `colorNames`, that assigns the closest value in the `colors()` list to any hexadecimal color specification.

We used these functions to assign names to the **Polychrome** 36 palette (Table 2).

**Table 2:** Color names assigned to the **Polychrome** 36 palette.

| ISCC | X11 |
| --- | --- |
| Dark_Purplish_Gray | gray33 |
| Purplish_White | gray89 |
| Vivid_Red | firebrick1 |
| Vivid_Purple | magenta |
| Vivid_Yellowish_Green | green |
| Strong_Purplish_Blue | dodgerblue |
| Vivid_Orange_Yellow | darkgoldenrod1 |
| Vivid_Purplish_Red | violetred3 |
| Brilliant_Green | aquamarine |
| Vivid_Yellow_Green | darkolivegreen3 |
| Vivid_Blue | lightskyblue |
| Brilliant_Purple | plum1 |
| Vivid_Violet | purple |
| Strong_Pink | lightpink1 |
| Strong_Blue | dodgerblue4 |
| Strong_Reddish_Orange | tomato3 |
| Vivid_Green | seagreen |
| Light_Olive_Brown | goldenrod4 |
| Vivid_Reddish_Purple | darkmagenta |
| Vivid_Greenish_Yellow | gold |
| Vivid_Yellowish_Green | springgreen3 |
| Vivid_Red | deeppink |
| Vivid_Purplish_Red | maroon1 |
| Pale_Yellow | navajowhite |
| Strong_Reddish_Purple | plum3 |
| Vivid_Violet | purple3 |
| Vivid_Yellow_Green | greenyellow |
| Very_Light_Blue | lightsteelblue1 |
| Strong_Reddish_Brown | tomato4 |
| Very_Light_Yellowish_Green | darkseagreen2 |
| Very_Light_Bluish_Green | darkslategray3 |
| Deep_Greenish_Blue | turquoise4 |
| Vivid_Purple | mediumorchid1 |
| Deep_Purple | mediumorchid4 |
| Brilliant_Blue | steelblue1 |
| Vivid_Violet | blue |

~~~
*R> names36 <− data.frame(ISCC = isccNames(P36), X11 = colorNames(P36))*
~~~

Interestingly, the ISCC set of names for the colors in this palette contain several duplicated names, including one—Vivid-Violet—that is used three times. To understand why, we produced a set of scatter plots that show the parts of U-V chromatic space used by different palettes and different naming systems (Figure 11). The “Polychrome 200” palette was created for this plot in order to give an idea of the “viewable” region of U-V coordinates; the other color sets are plotted so the coordinates of the axes match this one. The Polychrome 36 palette uses most of the available U-V-space. The set of UNIX/X11 color names also extend pretty much to the limits of visible U-V-space. However, there are some obvious gaps related to the fact that it is concentrated on radial vectors of constant hue. The ISCC-NBS color names, by contrast, are clearly more concentrated toward the center of U-V-space away from the more “vivid” colors near the boundary. This concentration may reflect a belief that vivid colors are harder to distinguish even if they are well-separated in CIE L*u*v* space.

**Figure 11:**
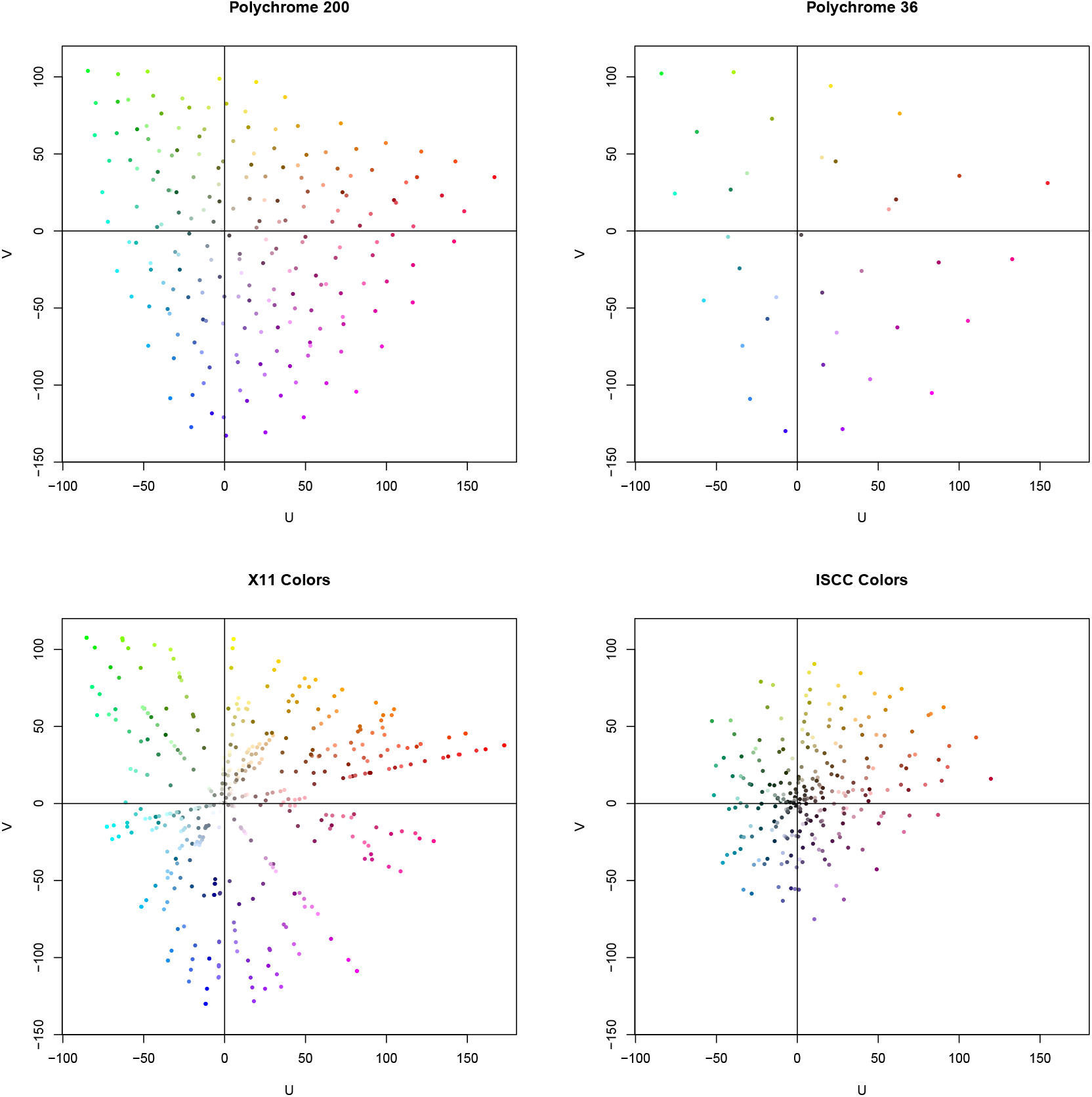
Scatter plot in the U-V chromatic coordinates of CIE L*u*v* space of different sets of colors. (Top Left) The **Polychrome** 200 palette; a set of 200 colors chosen to maximize their separation. (Top Right) The **Polychrome** 36 palette. (Bottom Left) Colors identified by names in the UNIX/X11 color list. (Bottom Right) Colors represented by name in the ISCC-NBS standard set of 267 colors.

## 6. A palette for spectral karyotyping

We now turn to the problem of constructing a palette for displaying data based on its chromosome of origin. As our starting point, we take a palette that is already in common use in spectral karyotyping. We downloaded electronic versions of the figures from two publications (Anguiano, Wang, Wang, Boyar, Mahon, El Naggar, Kohn, Haddadin, Sulcova, Sbeiti, Ayad, White, and Strom 2012; Imataka and Arisaka 2012)) that used hardware and software from Applied Spectral Imaging in Carlsbad, CA. The palettes in these images were manually extracted from the figures using the “eyedropper” tool in the GNU Image Manipulation Program (Peck 2006), and then saved in a CSV file that is included as extra data that accompanies the **Polychrome** package.

~~~
*R> oldsky <− read.csv(system.file(“extData/sky.csv”, package = “Polychrome”))
R> chr <− paste(“chr”, oldsky$Chromosome, sep = ‘‘)
R> temp <− t(sapply(oldsky$RGB4, function(x) {
+ eval(parse(text = paste(“c”, x, sep = ““)))
+ }))
R> dimnames(temp) <− list(chr, c(“R”, “G”, “B”))
R> oldsky.colors <− hex(sky.rgb <− RGB(temp/255))
R> rm(chr, temp, oldsky)*
~~~

This existing palette is displayed as the top panel of Figure 12. Some of the colors in this palette are difficult to distinguish, such as the greens of Chromosome 10 and Chromosome X, or the ligher shades in Chromosomes 3, 7, 9, 21, and 22. This difficulty is confirmed when we compute the minimum distances in CIE L*u*v* space, which suggests that only about 15 different colors are well-separated.

**Figure 12:**
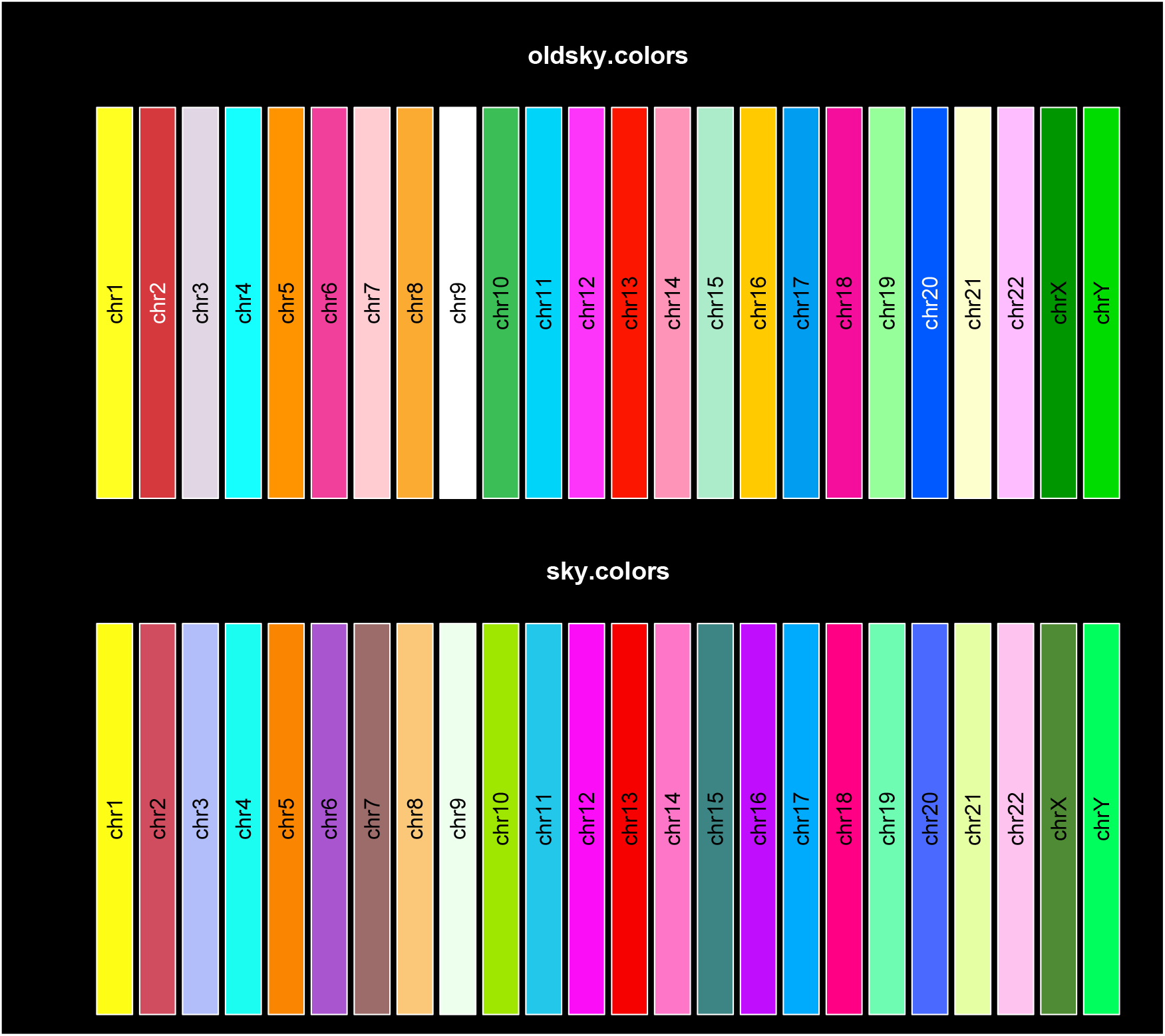
Palettes for displaying data based on the chromosome of origin. (Top) A 24-color palette in current use to display spectral karyotypes. (Bottom) The 24-color sky palette produced using **Polychrome**.

~~~
*R> od <− computeDistances(oldsky.colors)
R> min(od)*
~~~

~~~
[1] 15.54639
~~~

~~~
*R> sum(od > 39)*
~~~

~~~
[1] 15
~~~

We then used the `createPalette` function to construct an alternative palette. (See Appendix for details.) We seeded the algorithm using the two most distinguishable colors from the existing spectral karyotyping palette (the yellow of Chromosome 1 and the blue of Chromosome 20). Because these data are conventionally displayed on a black background, we restricted the luminance range to lie between 50 and 100. Because the original SKY colors were chosen in part to reflect the combination of fluorochromes used to label different chromosomes, we reordered our new palette to give the best matches to the older one. The resulting palette is shown in the bottom panel of Figure 12. The minimum separation between colors in this alternative palette is about 43 units.

~~~
*R> ns <− computeDistances(sky.colors)
R> min(ns)*
~~~

~~~
[1] 42.90331
~~~

~~~
*R> opar <− invertColors()
R> par(mfrow = c(2, 1))
R> swatch(oldsky.colors)
R> swatch(sky.colors)
R> par(opar)*
~~~

## 7. Displaying data by chromosome

In this final section, we will illustrate the application of **Polychrome** to the display of data that maps naturally to the human genome. In his thesis, Abrams developed a tool that converts karyotypes from text strings written using the International System for Human Cytogenetic Nomenclature (ISCN) into computationally friendly binary vectors, using three binary bits per cytoband (Abrams, Peabody, Heerema, and Payne 2015). We applied this tool to all 10,619 karyotypes from patients with acute myeloid leukemia (AML) available in the Mitelman Database of Chromosome Aberrations and Gene Fusions in Cancer (http://cgap.nci.nih.gov/Chromosomes/Mitelman). We then performed a principal components analysis (PCA) on the binary vectors to identify cytogenetically defined subtypes of AML.

In order to understand which factors were most strongly driving the PCA, we plotted the loadings associated with each cytoband (Figure 13). For the plot shown here, we have manually added labels that identify one cluster of points from each chromosome, selecting clusters that contribute most significantly to one of the first two principal components. The plot clearly shows that cytobands on the same chromosome tend to cluster together. Moreover, in many cases, there are at least two different clusters coming from each chromosome. For example, there is another (unlabeled) “midnight blue” cluster of data from chromosome 2 in the upper right of the figure, and a second (unlabeled) cluster of “fire brick” data from chromosome 1 at about (0.035, −0.015). In between the labeled clusters from chromosomes 9 and 22, there is also an unlabeled “dark goldenrod” set of data from chromosome 18. The observation that cytobands from the same chromosome tend to form two clusters results from the biological facts that many abnormalities in the Mitelman database occur on the scale of an entire chromosomal arm, and (most) chromosomes have two arms. (Note: the color names used in this paragraph were obtained by applying the `colorNames` function to the palette used to display the data.)

**Figure 13:**
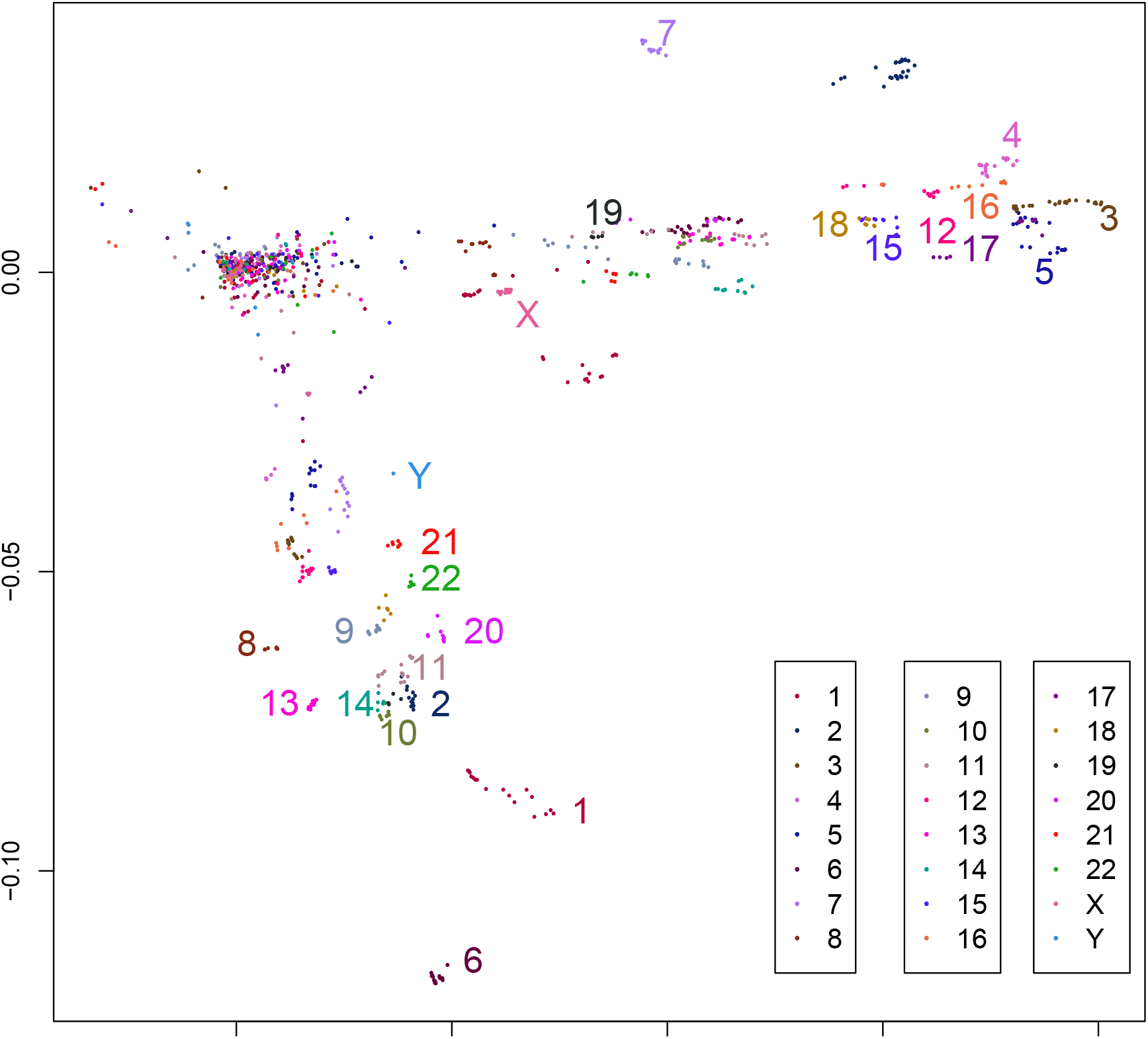
Plot of the loadings of cytoband data on the first two principal components (PC) of cytogenetic data on AML patients.

## 8. Conclusion

In this paper, we have reviewed many of the existing large color palettes available in R or in the published literature. Of these existing palettes, only the one introduced in Glasbey *et al.* (2007) contains at least 25 colors that are expected to be easily distinguished according to the criterion developed by Carter and Carter (1981, 1982).

To address this problem, we developed the **Polychrome** package, which provides tools to create palettes using an algorithm closely related to the one described by Glasbey *et al.* (2007). This is the first tool of its kind for R. We have demonstrated four different palettes (Polychrome 36, Dark 24, Light 24, and Sky), each of which contains at least 24 colors whose minimum separation is at least 40, thus meeting the Carter & Carter criterion. One potential weakness of the palettes produced here is that they rely on CIE L*u*v* space, which is designed to work well with light-based color systems like computer monitors. Some of the colors may not be as easy to distinguish when viewed using pigment-based colors. For example, colors NC6 and NC23 in the Dark 24 palette can be hard to distinguish on printed copies of the manuscript. If printing is the primary output medium, it might be better to use algorithms based on CIE L*a*b* space. The difference might also explain why some of the Glasbey colors were harder to separate in CIE L*u*v* space. We have also described additional tools to visualize and to assign meaningful names to palettes. We believe these palettes, and the tools to work with them, will be useful to all researchers who must display categorical data with a large number of factors.

## 9. Appendix

Here is the code used to create Figure 11

~~~
*R> library(“colorspace”)
R> cs <− hex(sRGB(t(col2rgb(colors()))/255))
R> names(cs) <− colors()
R> set.seed(451226)
R> p200 <− createPalette(200, c(“#ffffff”, “#010101”))
R> data(“iscc”)
R> opar <− par(mfrow = c(2, 2), mai = c(0.82, 0.82, 0.82, 0.42))
R> uvscatter(p200, xlim = c(−90, 170), ylim = c(−140, 110),
+ main = “Polychrome 200”)
R> abline(h = 0, v = 0)
R> uvscatter(P36, xlim = c(−90, 170), ylim = c(−140, 110),
+ main = “Polychrome 36”)
R> abline(h = 0, v = 0)
R> uvscatter(cs, xlim = c(−90, 170), ylim = c(−140, 110),
+ main = “X11 Colors”)
R> abline(h = 0, v = 0)
R> uvscatter(iscc$Hex, xlim = c(−90, 170), ylim = c(−140, 110),
+ main = “ISCC Colors”)
R> abline(h = 0, v = 0)
R> par(opar)*
~~~

Here is the code used to create the new spectral karyotping (SKY) color palette.

~~~
*R> sky.luv <− as(sky.rgb, “LUV”)
R> color.seed <− c(“#FFFF22”, “#0058FF”) # first two oldsky.colors R> set.seed(362425)
R> my.colors <− rev(createPalette(24, color.seed, range = c(50, 100),
+ M = 100000))
R> names(my.colors) <− names(oldsky.colors)
R> sky.colors <− my.colors R> sky.mat <− sky.luv@coords
R> my.mat <− as(hex2RGB(my.colors), “LUV”)@coords R> dmat <− matrix(NA, 24, 24)
R> for (i in 1:24) {
+ ed <− sqrt(apply(sweep(my.mat, 2, sky.mat[i,], “-”)^2, 1, sum))
+ dmat[i,] <− ed + }
R> rownames(dmat) <− names(oldsky.colors)
R> mnames <− colnames(dmat) <− sub(“chr”, “M”, names(oldsky.colors))
R> mapping <− matrix(NA, 24,2)
R> for (i in 1:23) {
+ mind <− apply(dmat, 1, min)
+ w <− which.min(mind)
+ v <− which(dmat[w,] == mind[w])
+ mapping[i,] <− c(names(w), names(v))
+ dmat <− dmat[−w,, drop = FALSE]
+ dmat <− dmat[, −v]
+ }
R> mapping[24,] <− c(names(oldsky.colors)[which(!(names(oldsky.colors)
+ %in% mapping[,1]))], mnames[which(!(mnames %in% mapping[,2]))])
R> foo <− as.numeric(sub(“Y”, 24, sub(“X”, “23”, sub(“M”, ““,
+ mapping[,2]))))
R> mapping <− mapping[order(foo),]
R> names(sky.colors) <− mapping[,1]
R> sky.colors <− sky.colors[names(oldsky.colors)]*
~~~

Here is the code to create the coor palette used for displaying data from the Mitleman database.

~~~
*R> chrColors <− rev(Dark24)
R> names(chrColors) <− paste(“chr”, c(1:22, “X”, “Y”))
R> amlPC <− read.csv(system.file(“extData/amlPC.csv”,
+ package = “Polychrome”),row.names = 1)
R> makeleg <− function(x, y, dx, dy = 0, chrColors, pch, …) {
+ lab <− c(1:22, “X”, “Y”)
+ legend(x, y, lab[1:8], col = chrColors[1:8], pch = pch, …)
+ legend(x + dx, y + dy, lab[9:16], col = chrColors[9:16],
+ pch = pch, …)
+ legend(x + 2*dx, y + 2*dy, lab[17:24], col = chrColors[17:24],
+ pch = pch, …)
+ }*
~~~

Finally, here is the code to produce Figure 13.

~~~
*R> plot (amlPC[, 2], amlPC[,3],
+ pch = 16, col=chrColors[amlPC[,1]],
+ xlab = “Loadings, PC1”, ylab = “Loadings, PC2”)
R> makeleg(0.05, −0.065, 0.012, 0, chrColors, 16, cex =1.2)
R> x <− c(0.032, 0.019, 0.081, 0.072, 0.075, 0.022, 0.04, 0.001,
+ 0.01, 0.015, 0.018, 0.065, 0.004, 0.011, 0.06,
+ 0.069, 0.069, 0.055, 0.034, 0.023, 0.019, 0.019, 0.027, 0.017)
R> y <− c(−0.09, −0.072, 0.009, 0.023, 0, −0.115, 0.04, −0.062,
+ −0.06, −0.077, −0.066, 0.007, −0.072, −0.072, 0.004,
+ 0.011, 0.004, 0.008, 0.010, −0.06, −0.045, −0.051, −0.007, −0.034)
R> ch <− c(1:22, “X”, “Y”)
R> cl <− c(1:24)
R> text(x, y, ch, col=chrColors[cl], cex = 1.5)*
~~~

